# Model-based deconvolution of a force signal to estimate motor unit twitch parameters under low, moderate and high force isometric contractions

**DOI:** 10.1101/2024.05.14.594072

**Authors:** Robin Rohlén, Jan Celichowski

## Abstract

Muscle force generation and human movement are organised by the central nervous system and executed by the peripheral nervous system and the muscle fibres through molecular and electrical mechanisms. Over the last half-century, attempts have been made to elucidate these mechanisms in vivo, primarily focusing on the motor unit (MU) activity because of its role as the smallest voluntarily contractible unit. Although it is firmly established that the nervous system controls muscle force by modulating MU activity, it is yet possible to distinguish between the activities of slow- and fast-twitch MUs non-invasively, which is important for rehabilitation and diagnostic purposes. Although different methods exist to extract MU twitch parameters from a force signal, no method can accurately identify a single MU twitch given a single MU spike train. We addressed this problem by developing a model-based deconvolution method. We evaluated the method using a MU-based recruitment model under isometric contractions and tested it on experimental data. We found that the deconvolution method can provide non-biased average twitch parameter estimates with low variance for the latest recruited MUs, irrespective of contraction level. It can estimate average twitch parameters when the underlying MUs comprise unequal successive twitch profiles, the force signal has lower signal-to-noise ratios, or when the spike train includes missed firings at the cost of slightly increased bias or variance. Finally, the method provides twitch parameter estimates that align with the expected MU recruitment characteristics in experimental conditions. To conclude, the deconvolution method may be used to study slow and fast MUs for rehabilitation and neuromuscular diagnostics.

**Author Summary:** To generate force voluntarily with a specific muscle, the brain plans and sends signals through the spinal cord via motor neurons, each of which communicates with a set of muscle fibres. Together, these muscle fibres and the motor neuron are called a motor unit. In the literature, the neural signals have received much attention, whereas the mechanical force-generating muscle fibres have received much less due to the limitations of current methods. By extracting the mechanical characteristics of these muscle fibres connected to a specific motor neuron type in vivo, one can use this information for rehabilitation and neuromuscular diagnostics of humans. Here, we proposed a method that can accurately estimate the force profile from each motor unit during low to high contraction levels. This method can be used for rehabilitation and neuromuscular diagnostics purposes.

## Introduction

Muscle force generation and human movement are organised by the central nervous system and executed by the peripheral nervous system and the muscle fibres through molecular and electrical mechanisms. Over the last half-century, attempts have been made to elucidate these mechanisms in animals and humans in vivo, primarily focusing on the motor unit (MU) activity because of its role as the smallest voluntarily contractible unit, each innervated by a single motor neuron. Although it is firmly established that the nervous system controls muscle force by modulating MU activity (1), it is yet possible to distinguish between the activities of slow- and fast-twitch MUs non-invasively in humans (2). Interestingly, the MU twitches’ contractile features, such as contraction time and maximal twitch amplitude, seem to be a useful diagnostic marker, as shown in a mouse model of amyotrophic lateral sclerosis due to the loss of fast-fatigable MUs (3). If we could extract MU twitch properties from slow and fast twitch MUs non-invasively in humans, it can be used for diagnostic purposes.

Methodologically, the study of MU twitches mostly relies on trains of *electrical* or *voluntary*-generated stimulus and directly recording the generated force pulling on the tendon with a force transducer. Given the recorded force signal of the partially fused tetanus, a computational method extracts the MU twitch parameters describing the contractile properties of each MU (4). Although there are different computational methods for this purpose in the literature, they are all subject to various limitations, which will be presented next.

Spike-triggered averaging (STA) is the most well-known computational method for extracting twitch parameter estimates for an average twitch. This method uses the time instants of the stimulus as triggers in the muscle force signal within a pre-defined window length and averages all the triggered windowed sub-signals. This method has been used in human studies, e.g., using intraneural motor-axon electrostimulation to obtain the stimulus used for averaging (5,6) and voluntary contractions using discharge times from high-density surface EMG (7) or intramuscular EMG (8–11).

Although STA has been widely used in the literature, it is well-known that it may provide biased twitch parameter estimates (12–14). The main reason for the resulting biasedness is that the time between two stimuli (inter-spike interval) is commonly 65-125 ms (8-15 Hz firing rate), which may be shorter than the contraction times that can be as long as 150 ms (15,16) and certainly shorter than the duration of the twitch (17). The partial fusion will lead to biased amplitude estimates as the triggers will capture the subtle oscillations of the partially fused tetanus (18).

A method overcoming such biases of the STA method is a model-based peel-off method (19,20). The method uses a parametrised twitch model fitted to the initial twitch in the force signal before subtraction, and the next twitch is being fitted and subtracted, etc. In this way, the method can correctly identify the twitch amplitudes, contraction times, and other twitch parameters in tetanic force signals (19,20) and tetanic displacement signals from ultrasound (21). This method can identify unequal successive twitch shapes, i.e., a twitch profile shape variation within the same MU, an important feature in force generation (22). On the other hand, if the signal-to-noise ratio is low, there may be a risk of error propagation if the preceding twitches are poorly estimated, and the error remains when estimating the consecutive twitch(es). Moreover, this method works only when a single MU is activated, likely only in cases stimulating a single axon in a specific rat model (23) or humans using microelectrodes (5). Although it is possible to electrically stimulate multiple axons simultaneously, even with irregular time intervals, to mimic the characteristics of natural contractions (24), this method is not feasible for signals recorded under voluntary contractions when a pool of MUs are co-active.

Due to the various limitations of the previous methods, a model-based deconvolution of the muscle force signal in the time domain has been proposed (25). This model consists of a parametrised twitch model and an optimisation algorithm to estimate the MU twitch parameters for an average twitch by numerically solving a least mean square problem. In contrast to STA and the model-based peel-off method, this method requires *multiple stimuli as input*, such as spike trains from multiple MUs, resulting in the method’s output being parameters for *multiple average twitches*. Although this method fits well with the developments of decomposition of high-density surface EMG, which can decode the EMG signals into multiple MU spike trains (26), the method breaks down for a single MU spike train input. In some situations, only a single stimulus train (e.g., MU spike train from intramuscular EMG or electrical stimulation) is accessible, and it should suffice to estimate the parameters of an average twitch. Also, the method is limited to low contraction levels.

If we could develop a model-based deconvolution method that can take one stimulus input and provide accurate MU twitch parameters for a single average twitch, this method may replace the widely known STA method and not be dependent on having multiple MU spike trains for MU twitch parameter estimation. Moreover, since high-threshold MUs are expected to contribute the most to the force signal (27), it should be possible to extract these twitches because of the relative amplitude differences between high-threshold and low-threshold MUs.

In this study, we aimed to develop a model-based deconvolution method to deconvolve the force-based unfused tetanus from an isometric voluntary contraction that can accurately estimate the MU twitch parameters for a single average twitch based on a single MU spike train. Besides evaluating the method using extensive simulations based on a recruitment and rate coding model (27) at various levels of synaptic input (from 2.5% to 70% of maximum synaptic input), we also compared it against the STA method. We further evaluated the effect of varying the twitch parameters within the same MU, the signal-to-noise levels of the force signal (from 20 dB to 0 dB) and the number of missed spikes in the spike train (from 0% to 5%). Finally, we tested the model-based deconvolution and STA methods on experimental recordings of the tibialis anterior muscle from 10% to 70% of the maximum voluntary isometric force to evaluate the methods’ feasibility based on the expected twitch characteristics.

## Methods

### Theory

The force generated by a muscle can be described as a sum of forces generated by a pool of MUs, where each MU comprises a train of firings or spikes (i.e., a spike train) and a twitch occurring after each spike (28). These spike-induced twitch profiles may vary across and within a MU (22). The generated force *Y*(*t*), *t* = [0, *T*] during an isometric contraction can be described by the following model:

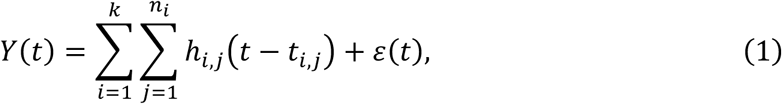

where *k* is the number of active MUs, *n*_*i*_ is the number of firings for the *i*th MU, *i* = 1, …, *k*, *h*_*i,j*_(*t* − *t*_*i,j*_) is the MU twitch response for the *j*th firing of the *i*th MU occurring at time instant *t*_*i,j*_, and *ɛ*(*t*) is additive noise. The spike train for the *i*th MU, containing zeros and ones, can be described as:

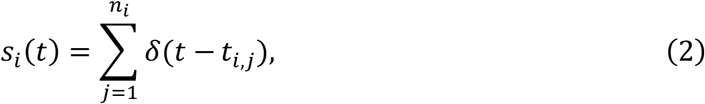

where *δ* is the Dirac Delta function.

Given a force signal, and *s*_*i*_(*t*), obtained from electrostimulation or estimated from EMG, the goal is to estimate the MU twitches to obtain the contractile characteristics of a MU. However, it is not feasible to estimate all the MU twitch profiles for a given MU *i* (i.e., *h*_*i,j*_, ∀*j* for a fixed *i*) unless only one active MU exists, such as in studies stimulating single axons in a rat model (23). Therefore, to obtain the contractile characteristics of a MU within a pool of active MUs, one must estimate a representative MU twitch, i.e., simplifying the model in Equation (1) by assuming the MU twitch responses to be identical within each MU, i.e., *h*_*i,j*_(*t*) = *h*_*i*_(*t*).

One alternative to a representative MU twitch is the average twitch. Replacing the varying twitch profiles with an average twitch (Fig. 1A) still captures the dynamics of the original force signal, although they are not identical (Fig. 1B). However, as stressed in numerous studies, STA leads to biased estimates of the average twitch (12–14). Instead, we implemented a model-based deconvolution method to provide unbiased estimates of a representative MU twitch.

**Fig 1.**
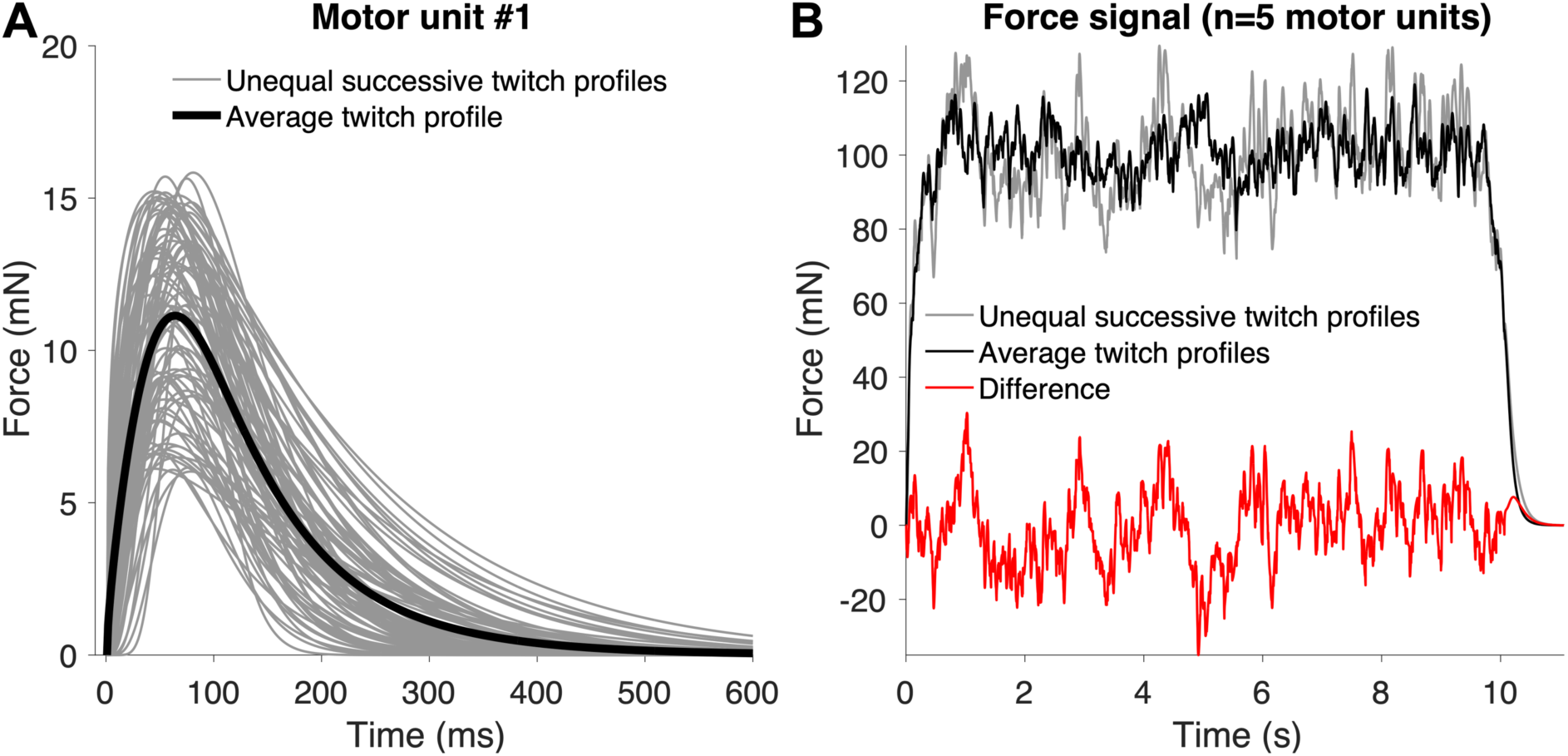
The difference in tetanic force using unequal successive MU twitch profiles and an average twitch profile using simulations. (**A**) Illustration of the varying MU twitch profiles (grey) and the average MU twitch profile (black). (**B**) Replacing the unequal successive twitch profiles with an average twitch (from A), the tetanic force (black) still captures the dynamics of the original force signal (grey), although they are not identical.

### Model-based deconvolution method

Inspired by the previous work of Negro and Orizio, who proposed a model-based deconvolution method with a *multi*-spike-train-input with a *multi*-twitch-parameter-output for low force levels (25), we implemented a model-based deconvolution method with a *single*-spike-train-input with a *single*-twitch-parameter-output. The method is based on using 1) an existing twitch model (20,29), 2) a high-pass filter, and 3) a particle swarm solver (30).

The twitch model has five parameters (*T*_*imp*_, *T*_*lead*_, *T*_*c*_, *T_hr_*, *F*_*max*_) corresponding to the time instant of the impulse (firing or spike), the time between spike and force onset, contraction time, half-relaxation time, and maximal twitch amplitude. Given its boundary conditions (Fig. 2A); the twitch for the *i*th MU can be described by the following equation (20):

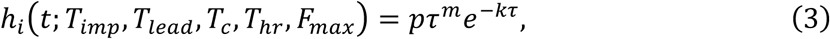

**Fig 2.**
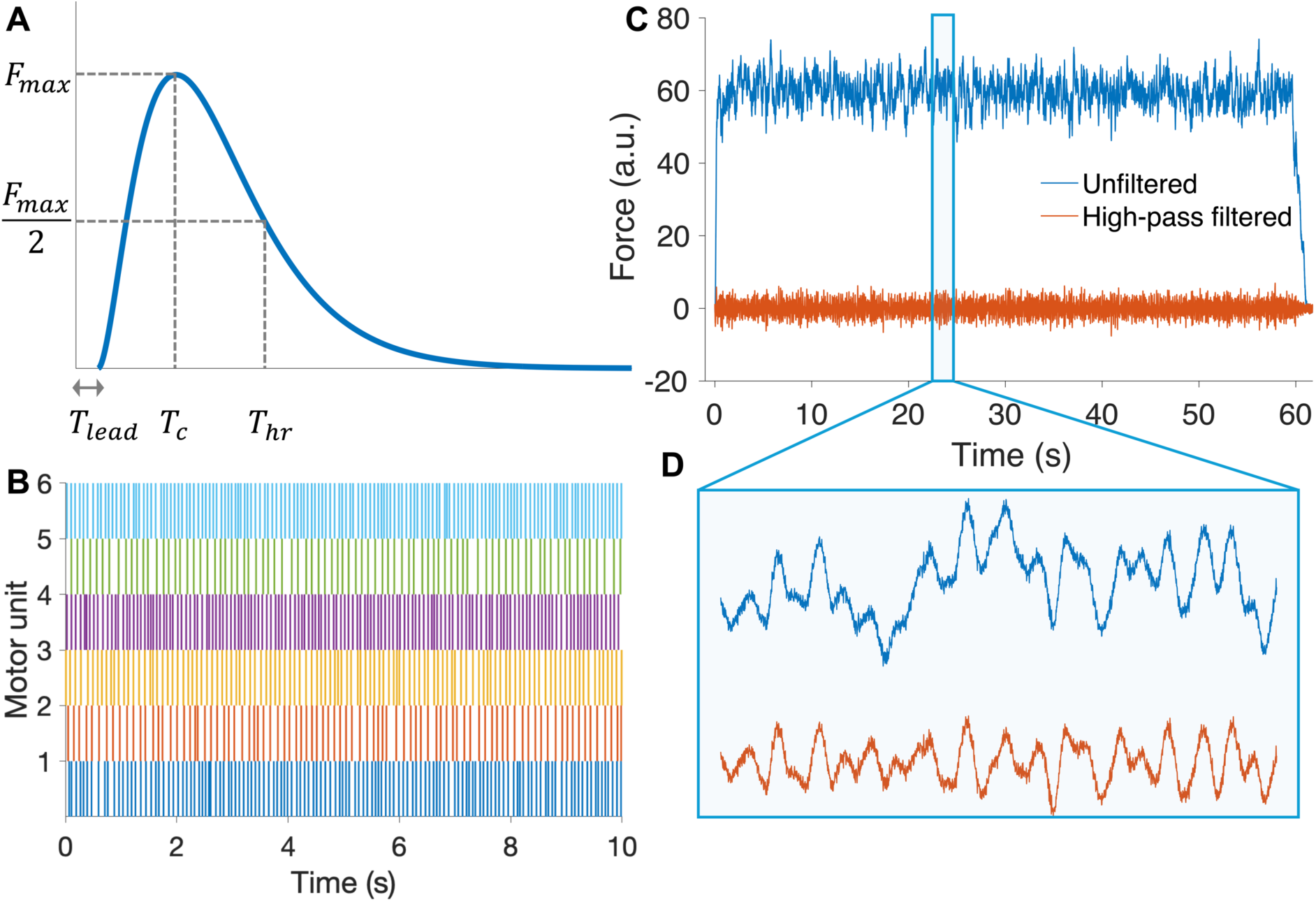
(**A**) The twitch model has five parameters (*T*_*imp*_, *T*_*lead*_, *T*_*c*_, *T*_*hr*_, *F*_*max*_) corresponding to the time instant of the impulse (firing or spike), the time between spike and force onset, contraction time, half-relaxation time, and maximal twitch amplitude. The time instant is zero in the figure. (**B**) The simulated motor unit (MU) spike trains and the simulated twitch profiles in (A) were used to obtain a tetanic force signal. (**C**) Integrating the twitch profiles over each MU spike train results in the simulated tetanic force signal (blue), a sum of forces generated by a pool of MUs. Given this tetanic force signal and a single MU spike train (B-C), estimating a MU twitch accurately using the existing model-based deconvolution method is difficult (25). The reason is that the tetanic signal of a single MU is much smaller than the (total) tetanic force signal, requiring an additional bias parameter intended to explain the remaining MU activities, and there are substantial amplitude modulations, making the task for the optimisation algorithm very complicated and easily converging in local minima. (**D**) We used a high pass filter to make the tetanic signal oscillate around zero and have less amplitude modulation, which makes the task easier for the optimisation algorithm to estimate the twitch parameters for a single MU.

where *τ* = *t* - *T*_*imp*_, *T*_*lead*_, 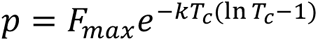, *m* = *KT_*c*_*, and 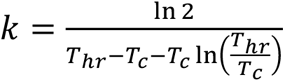. The time instants of the impulses (spike train) are usually assumed to be known through electrostimulation or estimated from EMG signal(s).

The sum of twitch profiles over each MU spike train (Fig. 2B) results in the tetanic force signal (Fig. 2C, blue line), which is a sum of forces generated by a pool of MUs. Given this tetanic force signal and a *single* MU spike train (Fig. 2B-C), it is difficult to estimate a MU twitch accurately using the existing model-based deconvolution method (25). The reason is that the tetanic signal of a single MU is much smaller in amplitude than the (total) tetanic force signal, requiring an additional bias parameter intended to explain the remaining MU activities, making the task for the solver very challenging and easily converging in local minima. To overcome this problem, we used a high-pass filter to make the tetanic signal oscillate around zero (Fig. 2D). Empirically, we found that a 4 Hz and 3^rd^ order high-pass Butterworth zero-phase filter provided the least biased and most stable performance for various simulation parameters (Fig. S1, Supporting Information). We also tested the effect of a low-pass filter, and we did not low-pass the signals in this work because it resulted in the least variance (Fig. S2, Supporting Information). Note that this may have to be considered on a case-by-case basis, depending on the noise level of the force signal.

The next step was to estimate the twitch parameters for a single MU (average) twitch based on the twitch model and the high-pass filter. We considered the following objective function to estimate the twitch parameters for the *i*th MU:

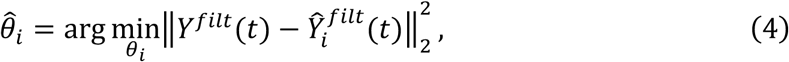

where *θ*_*i*_ = (*T*_*lead*_, *T*_*c*_, *T*_*hr*_, *F*_*max*_), *Y*^*filt*^(*t*) denote the high-pass filtered tetanic signal described in Equation (1), and 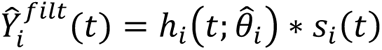 denote the high-pass filtered tetanic signal for the *i*th MU.

To estimate *θ*_*i*_ = (*T*_*lead*_, *T*_*c*_, *T*_*hr*_, *F*_*max*_) based on the objective function in Equation (4), *T*_*hr*_ was set to be a numerical ratio applied to the *T*_*c*_ value. This numerical ratio approach has been used in previous studies, e.g., 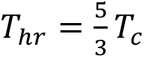 (25). We used a hybrid solver comprising a particle swarm solver (30), with a swarm size of 50, followed by a pattern search. The bounds of the parameters were set to be 0 ≤ *T*_*lead*_ ≤ 20 ms, 20 ≤ *T*_*c*_ ≤ 180 ms, 4 ≤ 3 × *T*_*hr*_/*T*_*c*_ ≤ 6, and 0.5 < *F*_*max*_ < 150 a.u. Since this study considered simulations to evaluate the method, we set the twitch amplitudes to be arbitrary units, which must be adapted to the scale of the experimental signals (see below).

### Spike-triggered averaging (STA) method

To compare the twitch parameter estimates from the model-based deconvolution method, we estimated the parameters using the STA method, i.e., taking the average of 200 ms windows for each MU spike as a trigger, resulting in a 200 ms STA twitch. Although it is well known that it may provide biased twitch parameter estimates (12–14), it has been widely used in the literature and still is considered the state-of-the-art method for a single MU spike train input. We estimated *T*_*c*_ by finding the time instant of the maximal amplitude of the STA twitch, and we estimated *F*_*max*_ by taking the difference between the maximum and minimum values of the STA twitch. The partial fusion will lead to biased amplitude estimates as the triggers will capture the subtle oscillations of the partially fused tetanus (18).

### Force simulations using a recruitment and rate coding model

To evaluate the performance of the model-based deconvolution and STA methods, we used a motor neuron-driven model with a recruitment and rate coding organisation (27) with a modified twitch response model (31). The simulations comprised *n* = 200 motor neurons where the time of the *i*th motor neuron’s *j*th spike was modelled as:

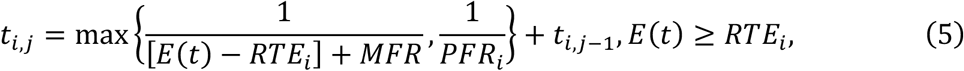

where *E*(*t*) is the excitatory drive representing the effective synaptic current at time *t*, *RTE*_*i*_ is the *i*th motor neuron’s recruitment threshold excitation (RTE), *MFR* is the minimum firing rate once a motoneuron was recruited (set to be 8 Hz), 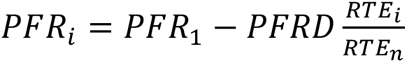 is the *i*th motor neuron’s maximal firing rate, which for the first motor neuron was set to be *PFR*_*1*_ = 35 Hz. The difference in peak firing rates between the 1^st^ and 200^th^ recruited motor neuron was *PFRD* = 10 Hz, as described in the original paper (27). RTE for the *i*th motor neuron was defined as 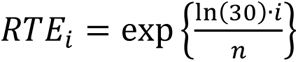, resulting in a range of recruitment threshold values (from 1 to 30). We imposed inter-spike interval (ISI) variability onto the model by adding a sample from a zero-mean normal distribution with a coefficient of variation equal to 15% for each spike.

The excitatory drive function *E*(*t*) was set to be a 60-s-long trapezoid function, i.e., a linear increase for the first 5 s from 0% to either 2.5, 30, or 70% of the maximum excitatory drive, and then a stable level of force for 50 s, and a linear 5 s decrease down to 0% (Fig. 3A). In Fig. 3, we illustrate the firing rate as a function of the excitatory drive (Fig. 3B).

**Fig 3.**
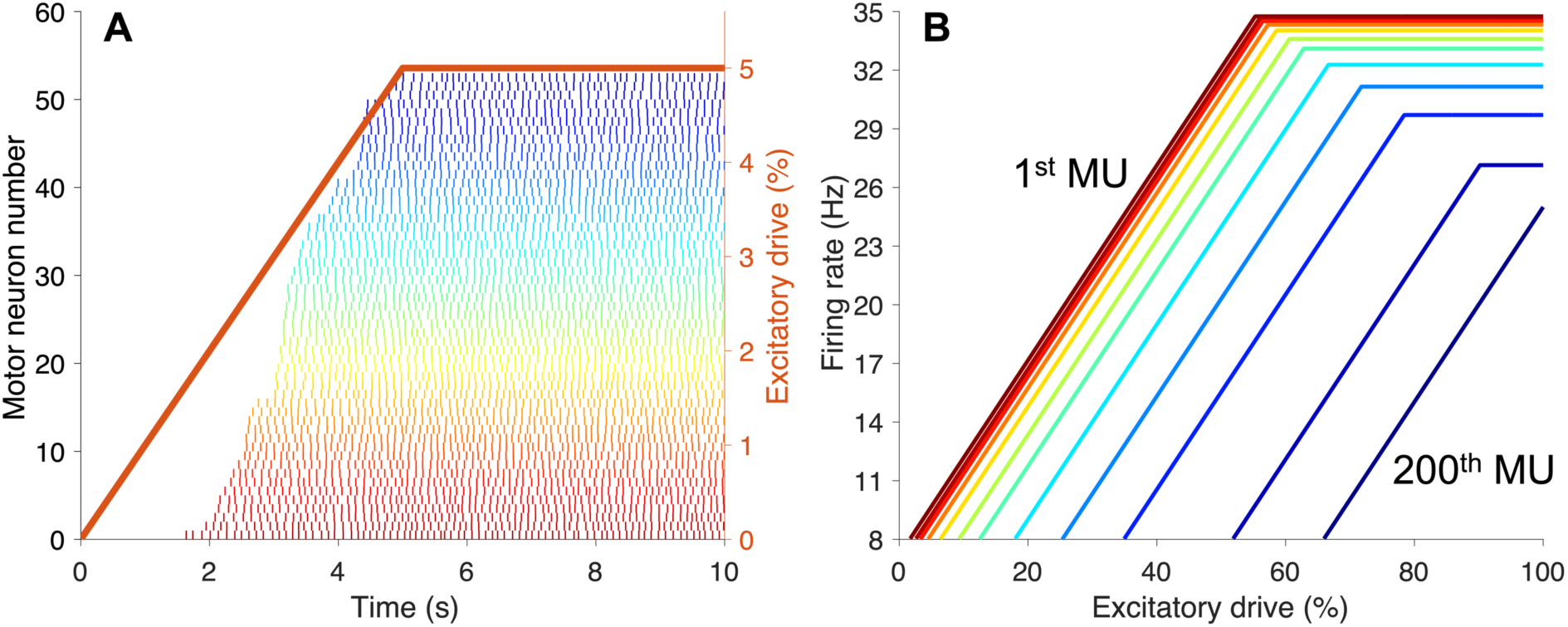
Motor unit (MU) recruitment model. (**A**) An example of the spike trains of recruited MUs given an excitatory drive function with 5% of the maximum excitatory drive. (**B**) The firing rate was set based on an excitatory drive function with an 8 Hz firing rate once any MU is recruited, and the maximum firing rate differs between MUs, with 35 Hz for the first and 25 Hz for the last (200th).

The force generated was achieved by summating a twitch profile for each spike and each MU as described in Equation 1. As described in a previous paper (31), the twitch profile parameters were generated orderly, representing slow and fast units. For instance, there was a 100-fold difference in twitch amplitude (*F*_*max*_) between the first and last MU in the pool (27). Therefore, the maximal twitch force for the *i*th MU was generated based on 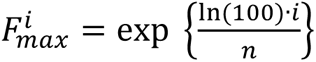, resulting in values from 1 to 100 a.u. (Fig. 4A). Similarly, the contraction time (*T*_*c*_) for each MU was generated based on an inverse power function depending on its maximal twitch force where we considered a 5-fold range in contraction time with the maximal contraction time being 150 ms (15,16) for the first recruited MU (Fig. 4A). Therefore, the contraction time for the *i*th MU was 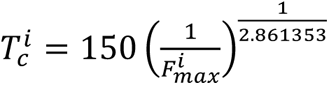. Similarly, the half relaxation time for the *i*th MU was 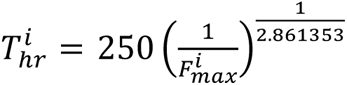 (Fig. 4A). This procedure resulted in one twitch profile for each MU having various shapes (Fig. 4B), i.e., equal successive twitch profiles within each MU.

**Fig 4.**
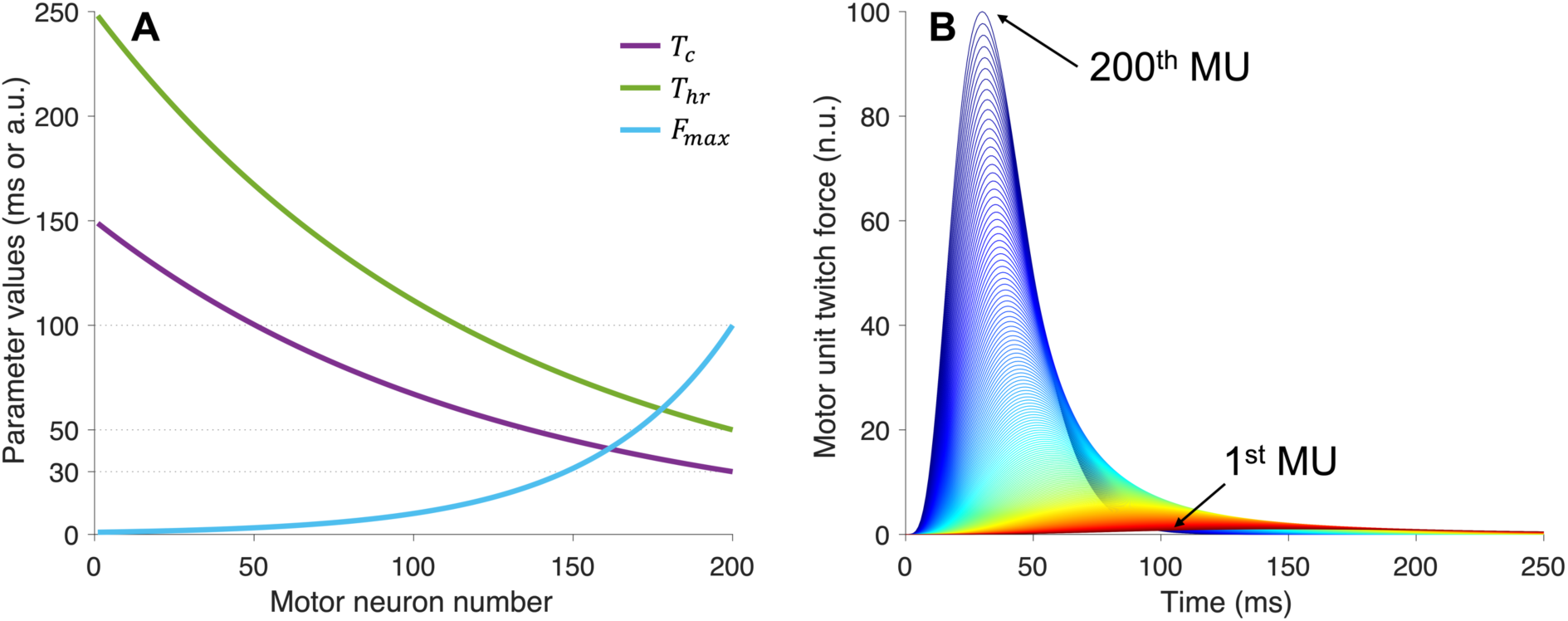
The twitch profile shape for a motor unit (MU) was generated based on three parameters: the contraction time (Tc), the half-relaxation time, and the maximum twitch force (Fmax). (**A**) Inverse power functions with ranges of values to mimic slow and fast MUs were used to generate the twitch parameters. (**B**) Illustration of the twitch profiles for the whole MU pool where the first recruited MUs had smaller twitch forces and longer contraction times than the latest recruited.

In physiological conditions, unequal successive twitch profiles better represent experimental data (22,31). Therefore, we also considered a simulation case with unequal successive twitch profiles within each MU (31) where the *i*th MU’s *j*th spike was randomly generated based on drawing values from 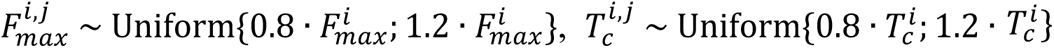, and 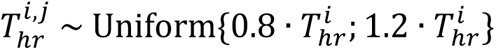.

Throughout all these scenarios, we randomly generated the spike-to-MU-twitch onset: *T*_*lead*_∼Uniform(0,15) ms. Finally, normally distributed white noise was added so that the signal-to-noise ratio was 20 dB. The sample rate was 1 kHz. Each case was repeated 20 times to increase the number of MU twitch parameter comparisons (ground truth vs estimated) for statistical means.

### Experimental data

To test and evaluate the feasibility of model-based deconvolution and STA methods for experimental recordings, a publicly available dataset from another study was included (32). The included data comprised eight subjects (27 ± 3 years old) with no history of lower limb injury or leg pain, performing isometric ankle dorsiflexions. EMG signals were recorded from the tibialis anterior muscle covered using four arrays of 64 surface electrodes (GR04MM1305, 13×5 gold-coated electrodes, with one absent, and a 4 mm inter-electrode distance; OT Bioelettronica, Italy) while simultaneously recording the generated force in an ankle dynamometer (OT Bioelettronica, Turin, Italy) with 60-s trapezoid curves as biofeedback. Before placing the electrodes, the skin was shaved and cleaned, and an adhesive foam and conductive paste were used to provide good skin-electrode contact. Reference electrodes were placed around the ankle. The data was sampled in monopolar derivation with 2048 Hz, amplified (x150), 10-500 Hz band-pass filtered, and digitised with a 16-bit resolution using the Quattrocento system (OT Bioelettronica, Turin, Italy). After performing two maximal voluntary contractions (MVC) the session consisted of eight isometric contractions performed at 10% to 80% of the MVC with 10% increments in a randomised order. After data collection, the signals were decomposed using a convolutive blind source separation method to identify MU spike trains, which were edited to improve accuracy. The details of the methodology are presented in the paper, providing the dataset (32).

Note that the publicly available data also included isometric knee extensions using the vastus lateralis, but we decided not to include these because the force recordings were not saved similarly to the tibialis anterior recordings, and they were of lower quality. After the initial inspection, three of the eight subjects from the tibialis anterior data had the same issue as in the vastus lateralis data, leading to omitting these three subjects. We omitted the 80% MVC recordings because they had fewer or no MUs. Finally, we only used data from one of the four grids based on the one with the most identified MUs because of the redundant information in the other grids and to limit the processing time.

### Statistical analysis

We tested whether the difference between the ground truth and estimated twitch parameters for each case, for each method described in the previous sections, significantly differed from zero with the nonparametric Sign Test. We also tested whether there was any difference in variation when the force signal had a lower signal-to-noise ratio (20 dB to 0 dB) and when there were missed spikes (0% to 5%), using the nonparametric Ansari-Bradley Test (after removing the median from each sample). We considered a significance level of 0.05.

## Results

For the equal successive twitch profile evaluation, the proposed model-based deconvolution method provided unbiased and least variance for the contraction time (*T*_*c*_), the half relaxation time (*T*_*hr*_), and the maximal twitch amplitude (*F*_*max*_) parameters of the latest recruited MUs at 2.5%, 30%, and 70% excitation maximum (Fig. 5B-D; Table S1, Supporting Information). Moreover, we found that the spike-to-force-onset (*T*_*lead*_) estimate had a high variance irrespective of the recruitment threshold (Fig. 5A), suggesting that it is better to fix this parameter before estimation in practice.

**Fig 5.**
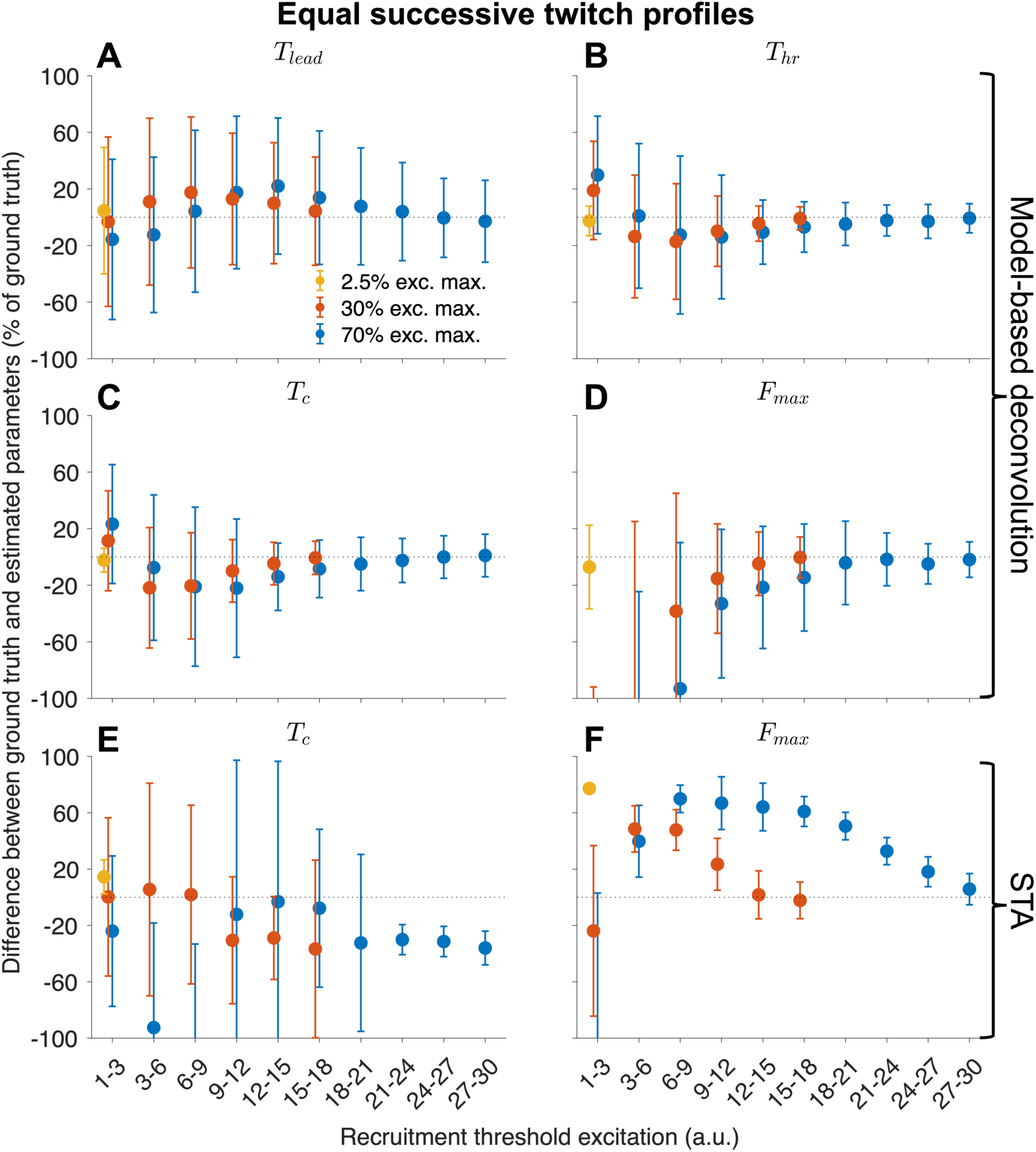
Twitch parameter estimation in the equal successive twitch profile case. (**A**) The spike-to-force-onset (Tlead) estimates using the model-based deconvolution method led to high variance irrespective of the recruitment threshold. (**B**) On the other hand, the half-relaxation time (Thr) for the latest recruited MUs at various excitatory drive functions could be reliably estimated with no bias and low variance. (**C**) The estimated contraction time (Tc) was very similar to the Thr shown in (B). (**D**) Similar to Tc and Thr, the maximal twitch amplitude (Fmax) could be reliably estimated for the latest recruited MUs. (**E**) The STA method provided biased contraction times. The variance differed depending on excitatory drive function and recruitment threshold. (**F**) Although the STA method provided relatively low variance in the Fmax estimates, they were, in most cases, biased in a non-linear way. However, it provides accurate Fmax estimates of the very latest recruited MUs, likely due to the low firing rates and lower contraction times of these MUs.

The STA method provided mostly biased twitch parameter estimates irrespective of recruitment threshold or excitatory drive function (Fig. 5E-F; Table S2, Supporting Information). Although the STA method provides fairly accurate maximal twitch amplitude (*F*_*max*_) estimates for the very latest recruited MUs (Fig. 5F), likely due to the lower firing rates and contraction times of these MUs, the model-based deconvolution method can estimate the twitch parameters of a larger MU pool than the STA method with the same or better precision (Fig. 5D; Tables S1-S2, Supporting Information).

When the force signal has a lower signal-to-noise ratio (from 20 dB to 0 dB), the model-based deconvolution method provides estimates with a higher variation (p < 0.001), although still unbiased (p > 0.05; Fig. S3, Supporting Information). Note that this was without low pass filtering the force signal. If the deconvolution method only uses 95% of the spikes (5% missed spikes), the estimate variation is the same, but there is an increased underestimation for the higher recruitment thresholds (p < 0.001; Fig. S4, Supporting Information).

In physiological conditions, unequal successive twitch profiles better represent experimental signals on MU level (22,31). Therefore, we also considered a simulation case with unequal successive twitch profiles within each MU. We found that the model-based deconvolution provided similar results to the equal successive twitch profile case with increased variance for the twitch parameters of the latest recruited MUs and a slight offset from the baseline (Fig. 5C-D vs. Fig. 6A-B). The STA method provided biased twitch parameter estimates where the contraction times had large variations (Fig. 6C-D).

**Fig 6.**
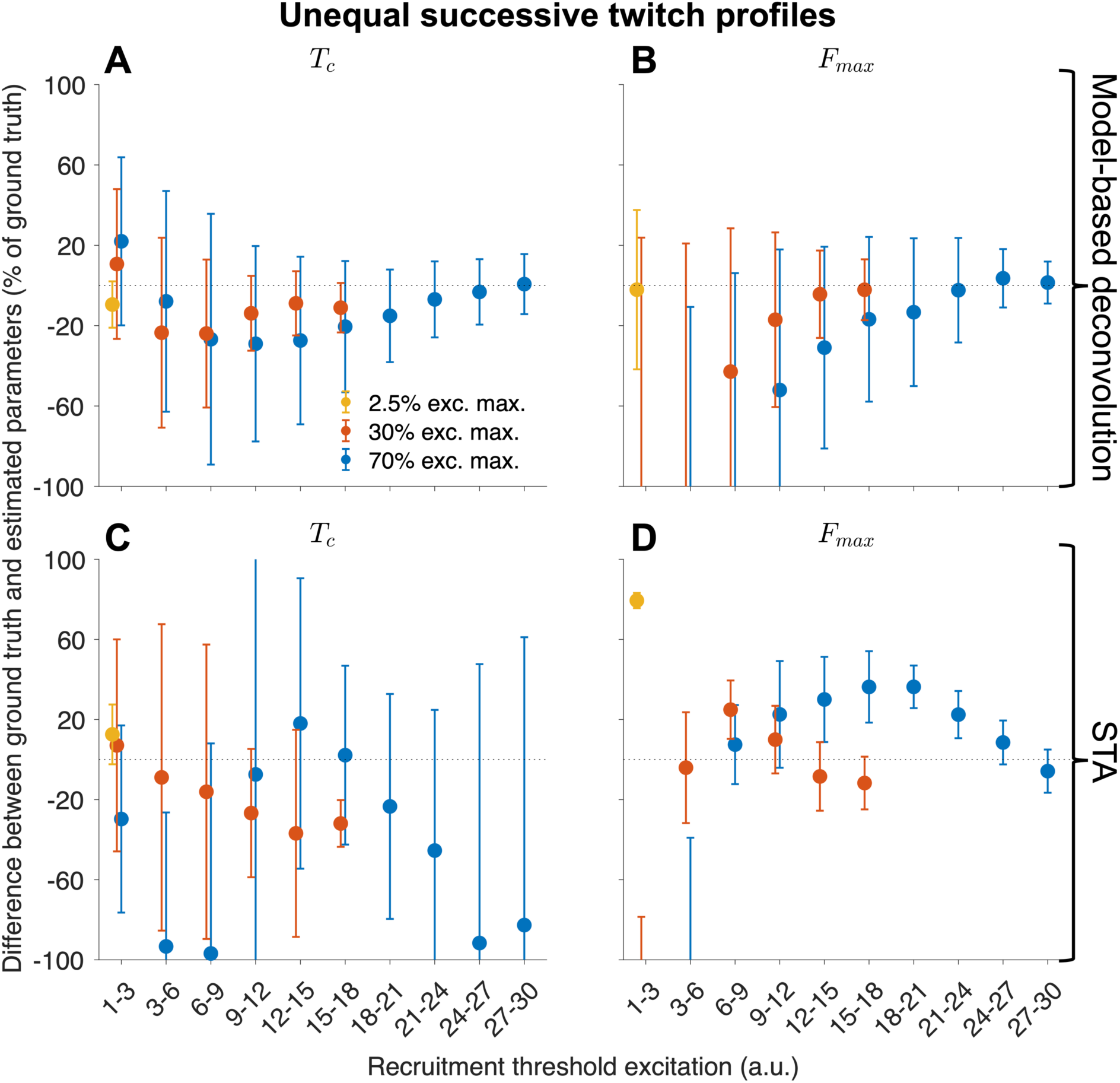
Twitch parameter estimation in the unequal successive twitch profile case. (**A**) The model-based deconvolution provided similar results to the equal successive twitch profile case, where the contraction times of the latest recruited motor units (MUs) at various excitatory drive functions could be reliably estimated with no bias and low variance. (**B**) Similar to Tc, the maximal twitch amplitude (Fmax) could be reliably estimated for the latest recruited MUs. Compared to the equal successive twitch profile case, there was slightly increased variance for the twitch parameters of the latest recruited MUs and a slight offset from the baseline. (**C**) Similar to the equal successive twitch profile case, the STA method provided biased contraction times. The variance differed depending on excitatory drive function and recruitment threshold. (**D**) In addition, the STA method provided relatively low variance in the Fmax estimates, and unlike the equal successive twitch profile case, the very latest recruited MUs were overestimated.

To test and evaluate the feasibility of model-based deconvolution and STA methods for experimental recordings, we estimated contraction times (*T*_*c*_) and maximal twitch amplitudes (*F*_*max*_) for 683 MUs in 10% MVC to 70% MVC (minimum of 76 MUs at 70% and maximum of 106 MUs at 30% MVC). In a representative case from one subject shown in Fig. 7, we show the contraction times (*T*_*c*_) estimated from the deconvolution method and how they decrease with increasing recruitment thresholds (Fig. 7A). Moreover, the maximal twitch amplitude (*F*_*max*_) increases with increasing recruitment thresholds (Fig. 7B). The STA method provides a less clear relationship regarding the contraction time (Fig. 7C), whereas the maximal twitch amplitude very clear but lower values compared to the deconvolution method (Fig. 7D), as seen in simulation cases (Figs. 5-6). These contraction times and maximal twitch amplitude relationships with the recruitment threshold are expected, as shown in the simulation model (Fig. 4A).

**Fig 7.**
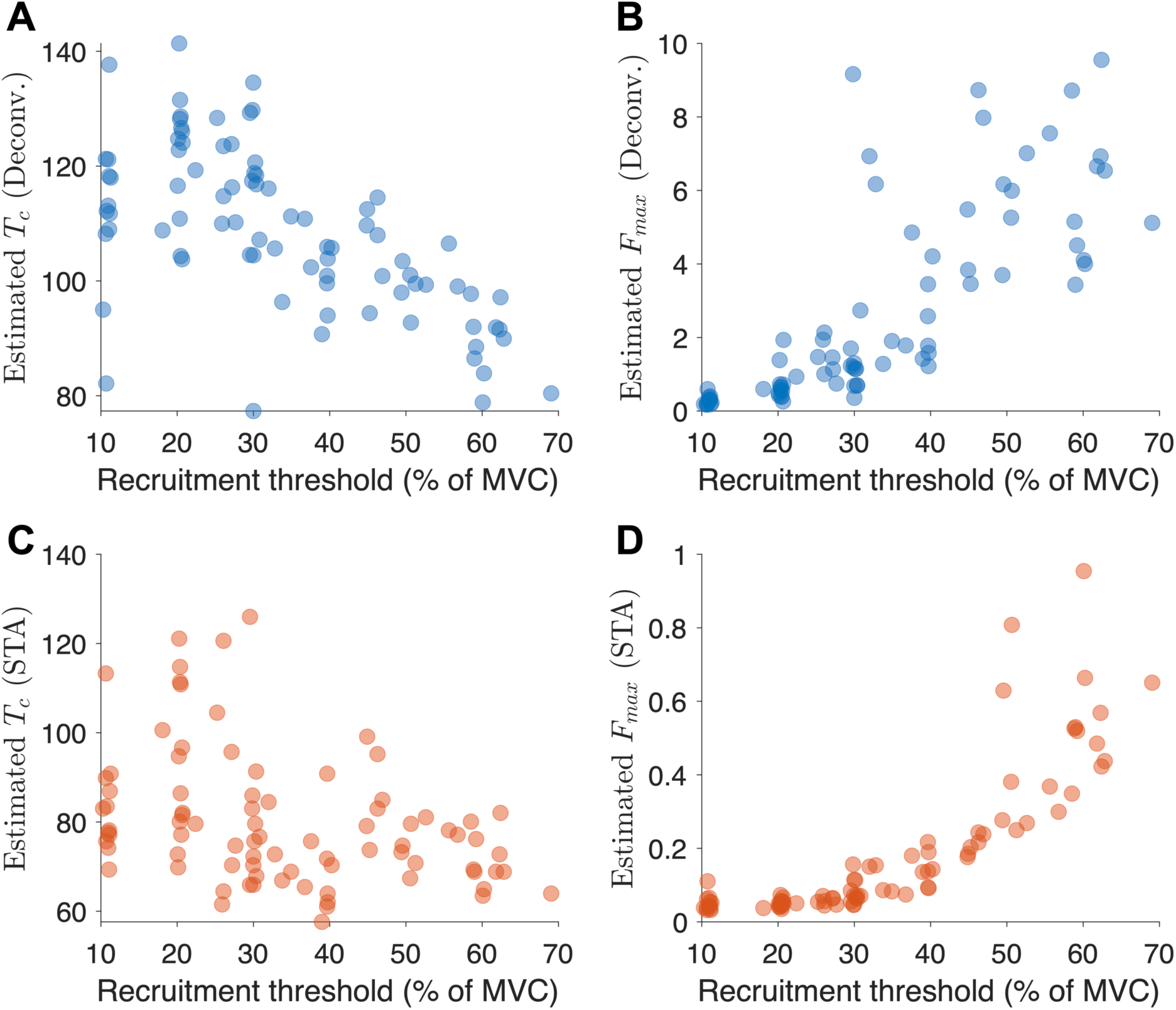
A representative case from a subject from experimental data comparing the estimated twitch parameters against the recruitment thresholds in terms of maximum voluntary contraction (MVC). (**A**) As expected, the contraction times (Tc) estimated using the model-based deconvolution method show a negative association between contraction time and recruitment threshold. (**B**) As expected, the maximal twitch amplitudes (Fmax) estimated using the model-based deconvolution method show a positive association between the twitch amplitude and recruitment threshold. (**C**) The spike-triggered averaging (STA) method shows a less clear relationship between the contraction time and recruitment threshold. (**D**) The STA method shows a clear positive association between the maximal twitch amplitude and the recruitment threshold.

By pooling all the estimated contraction times from the five subjects using the model-based deconvolution method, we observed a negative association with the recruitment threshold on the group level (Fig. 8A). Moreover, we observed a positive association between the estimated maximum twitch amplitude and the recruitment threshold (Fig. 8B). However, using the STA method, these relationships were less clear (Fig. 8C-D).

**Fig 8.**
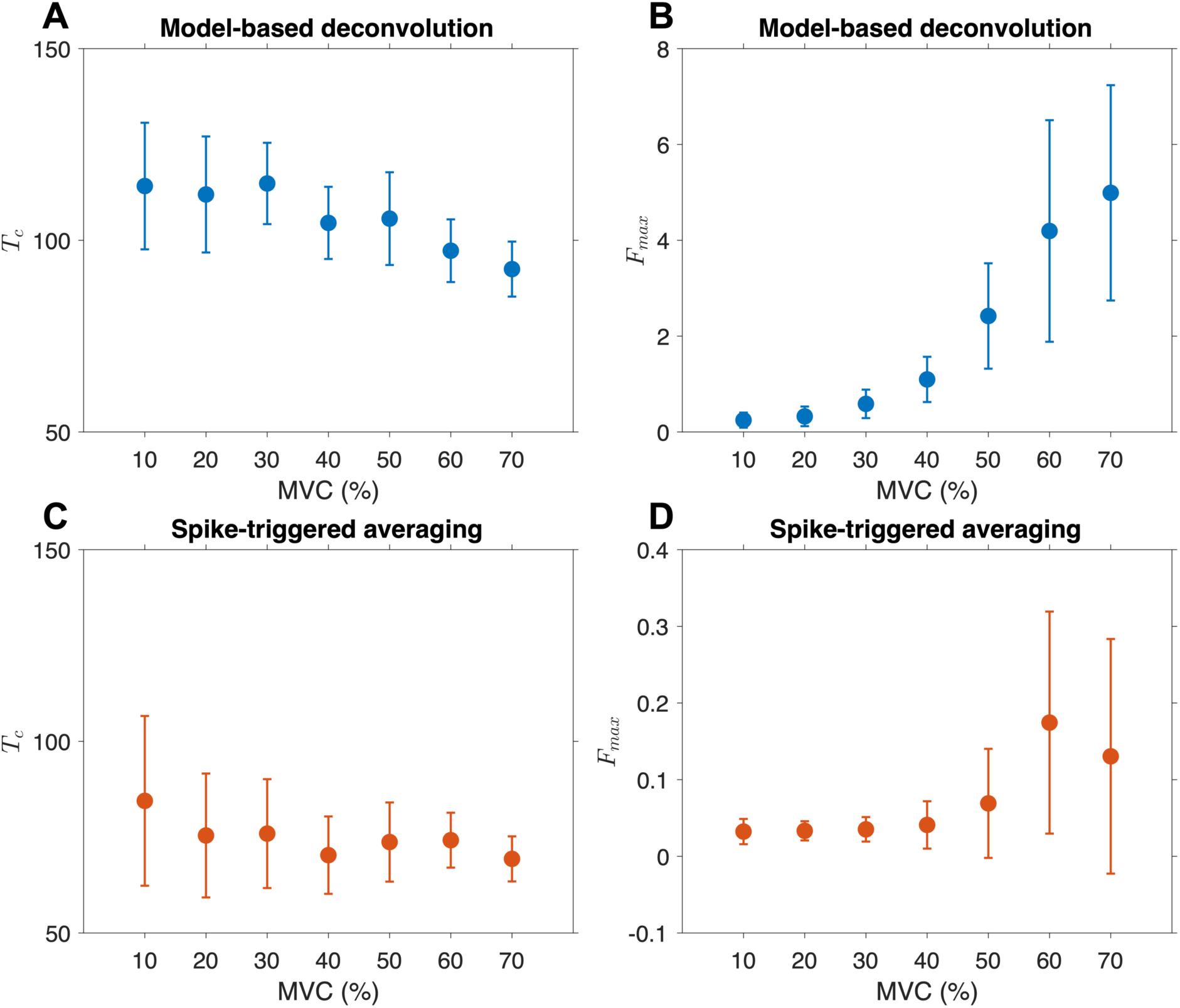
Pooling the estimated twitch parameters from all subjects from experimental data and comparing the estimated twitch parameters against the recruitment thresholds in terms of maximum voluntary contraction (MVC). (**A**) The contraction times (Tc) estimated using the model-based deconvolution method show a negative association between contraction time and recruitment threshold. (**B**) The maximal twitch amplitudes (Fmax) estimated using the model-based deconvolution method show a positive association between the twitch amplitude and recruitment threshold. (**C**) The spike-triggered averaging (STA) method shows a less clear relationship between the contraction time and recruitment threshold. (**D**) The STA method shows a less clear relationship between the maximal twitch amplitude and the recruitment threshold.

## Discussion

This study aimed to develop a model-based deconvolution method of a force-based signal of unfused tetanus that can accurately estimate the MU twitch parameters for a single average twitch based on a single MU spike train. The proposed method used a twitch model and an optimisation algorithm to estimate the twitch parameters from a high-pass filtered tetanic force signal. We performed extensive simulations using a recruitment and rate coding model to evaluate the method’s performance at low-, medium-, and high levels of synaptic input during isometric contractions. For comparison, we also used the STA method. Finally, we tested the model-based deconvolution and STA methods on experimental recordings of the tibialis anterior muscle from 10% to 70% of the maximum voluntary isometric force to evaluate the methods’ feasibility based on expected behaviour. Four main findings are presented and discussed below.

First, the model-based deconvolution method can provide non-biased average twitch parameter estimates with low variance for the latest recruited MUs, irrespective of contraction level. In other words, the method is biased towards higher twitch amplitudes and, therefore, later recruited ones according to Henneman’s size principle (33,34). One should stress that this is relevant for many methods interested in decoding compound signals into MU activity, including high-density surface EMG decomposition (35) and ultrafast ultrasound (31). Therefore, due to the relative differences in maximal twitch amplitude, the method cannot reliably estimate the twitch parameters of the early recruited MUs at medium-to-high contraction levels. To extract reliable twitch contraction times and maximum twitch amplitudes across a large population of the low- and high-threshold MU pool, one must record various contraction levels separately and extract the latest recruited ones (see 2.5%, 30%, and 70% of excitation maximum in Fig. 5C-D).

Second, the STA method provides biased twitch parameter estimates, with the contraction times generally overestimated and the maximal twitch amplitudes underestimated. The main reason for the resulting biasedness is that the inter-spike interval is commonly 65-125 ms (corresponding to an 8-15 Hz firing rate), which may be shorter than the contraction times that can be as long as 150 ms (15,16) and certainly shorter than the duration of the twitch (17). The partial fusion will lead to biased amplitude estimates as the triggers will capture the subtle oscillations of the partially fused tetanus (18). On the other hand, the maximal twitch amplitude values have low variation (see Figs. 5-6), suggesting that, together with information on firing rates, they can be used to adjust the bias. This remains to be investigated in future works. However, despite the biasedness, the STA method is simple and can be used initially as a sanity check before applying the model-based deconvolution method.

Third, the deconvolution method can estimate average twitch parameters when the underlying MUs comprise unequal successive twitch profiles, the force signal has lower signal-to-noise ratios, and the spike train includes missed firings at the cost of slightly increased bias or variance. Regarding the unequal successive twitch profiles, the method accurately estimates a representative twitch when the force signal comprises unequal successive MU twitch profiles within each MU. Although this representative varies around the average twitch, its variation is low enough to be a feasible representative twitch. Moreover, if the underlying unequal successive twitches are more similar, we should expect the estimated MU twitch to be closer to the average twitch. Regarding the lower signal-to-noise ratio of the force signal, there was an increase in the variance of the estimates. This may be overcome using low-pass filtering, which we did not explore in this study since low-pass filtering was not found useful in our baseline case (Fig. S2, Supporting Information). Regarding the missed spikes, there are various solutions to overcome the increased underestimation. For instance, one may adjust the parameters if it is possible to estimate the number of missed firings, e.g., during an isometric and stable contraction. In some situations, like electro-stimulations or using intramuscular EMG, we may be much more certain not to have any missed spikes. Taken together, unequal successive twitch profiles, lower signal-to-noise ratios, and when the spike train includes missed firings comes at the cost of slightly increased bias or variance, it is still within an acceptable range in practice with potential solutions to reduce or adjust for these issues.

Fourth, the deconvolution method provides twitch parameter estimates that align with the expected MU twitch characteristics, as shown in Fig. 4A. As summarised by Fuglevand and colleagues, the MU twitch parameters based on experimental findings show that 1) the twitch force over a pool has a wide range, 2) majority of MUs produce smaller forces and a few MUs generate high force, and 3) the force generated by a MU relates to the recruitment threshold (27). As shown in Figs. 7-8, this aligns with these three points. This finding shows the possibility of estimating MU twitch parameters across the MU pool, including fast MUs, suggesting the feasibility of using this method for rehabilitation and neuromuscular diagnostic purposes. For instance, it may track contractile function over time in reinnervated muscles. Moreover, the MU twitch parameters may be a diagnostic marker, as shown in a mouse model of amyotrophic lateral sclerosis, due to the loss of fast-fatigable MUs (3). Whether this method will play a role in rehabilitation and diagnostics remains to be seen.

The present study has three main limitations. *First*, the obtained twitches are representative twitches, likely close to the average twitch, whereas the twitches decomposed from the tetanic contractions of single MUs reveal some variability in amplitude and twitch time parameters (22). For most MUs, the single twitch force is lower than for twitches decomposed from unfused tetanus, and therefore, the proposed method may overestimate the twitch amplitude. *Second*, *T*_*lead*_ cannot be reliably estimated, likely due to changes in *T*_*c*_ and *T*_*hr*_ affecting the objective function more than changes in *T*_*lead*_. In previous studies with rat models, it turns out that the *T*_*lead*_ parameter values are very similar (19,20). So, an alternative solution would be to use previous knowledge or estimate *T*_*lead*_ before fixing it in the optimisation algorithm. For instance, the existing model-based deconvolution method using multiple spike trains as input assumed *T*_*lead*_ to be zero. Moreover, it is unclear how *T*_*lead*_ relates to the so-called neuromechanical delay that exists in the literature, which has been reported to be anything from 200 ms (36) to 500 ms (37), which is likely to be correlated with the contraction time as well as the *T*_*lead*_ parameter. *Third*, the simulation model was evaluated using a 5-fold difference in contraction times and half-relaxation times, and some muscles may have less difference, which may lead to identifying fewer accurate MU twitch parameters.

In conclusion, given a MU spike train, we have proposed a model-based deconvolution method to accurately decode a tetanic force signal at a low, medium, and high force level into a representative MU twitch. This method can replace the well-known STA method shown here, which provides biased twitch parameter estimates. The model-based deconvolution method may study slow and fast MUs for rehabilitation and neuromuscular diagnostics.

## Acknowledgements

The work was supported by funding from the Royal Physiographic Society of Lund. R.R. is supported by the Swedish Research Council (2023-06464), the Swedish Brain Foundation (PS2022-0021), the Swedish Research Council for Sport Science (FO2024-0003), and the Promobilia Foundation (A23161).

## Supporting information captions

***Fig S1.*** *Finding the optimal high-pass filter cutoff by comparing the performance between ground truth and estimated parameters for simulations with a trapezoid excitatory drive function with 2.5% of the maximal excitatory drive. 4 Hz high-pass filter cutoff provides the less biased estimates*.

***Fig S2.*** *Finding the optimal low-pass filter cutoff by comparing the performance between ground truth and estimated parameters for simulations with a trapezoid excitatory drive function with 2.5% of the maximal excitatory drive. Using no low-pass filter provides the least variation*.

***Fig S3.*** *When the force signal has a lower signal-to-noise ratio (from 20 dB to 0 dB), the model-based deconvolution method provides estimates with a higher variation (p < 0.001), although still unbiased (p > 0.05). Note that this was without low pass filtering the force signal*.

***Fig S4.*** *If the deconvolution method only uses 95% of the spikes (5% missed spikes), the estimate variation is the same, but there is an increased underestimation for the higher recruitment thresholds (p < 0.001)*.

***Table S1.*** *Nonparametric Sign Test for the difference between the ground truth and estimated twitch parameters from the model-based deconvolution method for equal and unequal successive twitch profile cases. *** p < 0.001, ** p < 0.01, * p < 0.05*.

***Table S2.*** *Nonparametric Sign Test for the difference between the ground truth and estimated twitch parameters from the spike-triggered averaging method for equal and unequal successive twitch profile cases. *** p < 0.001, ** p < 0.01, * p < 0.05*.

## Notes

### Competing Interest Statement

The authors have declared no competing interest.

### Summary of Updates

Evaluating the method with another simulation model at various contraction levels. Also included experimental data for testing the method.

## References

1. Farina D, Negro F. Common Synaptic Input to Motor Neurons, Motor Unit Synchronization, and Force Control. Exercise and Sport Sciences Reviews. 2015 Jan;43(1):23.

2. Ota K, Sasaki K. Influence of temperature on twitch potentiation following submaximal voluntary contractions in human plantar flexor muscles. Physiol Rep. 2023 Aug 24;11(16):e15802.

3. Chen X, Sanchez GN, Schnitzer MJ, Delp SL. Microendoscopy detects altered muscular contractile dynamics in a mouse model of amyotrophic lateral sclerosis. Sci Rep. 2020 Jan 16;10(1):457.

4. Raikova R, Krutki P, Celichowski J. Skeletal muscle models composed of motor units: A review. Journal of Electromyography and Kinesiology. 2023 Jun 1;70:102774.

5. McNulty PA, Macefield VG. Intraneural microstimulation of motor axons in the study of human single motor units. Muscle & Nerve. 2005;32(2):119–39.

6. Thomas CK. Human motor units studied by spike-triggered averaging and intraneural motor axon stimulation. Adv Exp Med Biol. 1995;384:147–60.

7. Cescon C, Gazzoni M, Gobbo M, Orizio C, Farina D. Non-invasive assessment of single motor unit mechanomyographic response and twitch force by spike-triggered averaging. Med Biol Eng Comput. 2004 Jul 1;42(4):496–501.

8. Milner-Brown HS, Stein RB, Yemm R. The contractile properties of human motor units during voluntary isometric contractions. The Journal of Physiology. 1973;228(2):285–306.

9. Nordstrom MA, Miles TS, Veale JL. Effect of motor unit firing pattern on twitches obtained by spike-triggered averaging. Muscle & Nerve: Official Journal of the American Association of Electrodiagnostic Medicine. 1989;12(7):556–67.

10. Roatta S, Arendt-Nielsen L, Farina D. Sympathetic-induced changes in discharge rate and spike-triggered average twitch torque of low-threshold motor units in humans. The Journal of Physiology. 2008;586(22):5561–74.

11. Stein RB, French AS, Mannard A, Yemm R. New methods for analysing motor function in man and animals. Brain Research. 1972 May 12;40(1):187–92.

12. Calancie B, Bawa P. Limitations of the spike-triggered averaging technique. Muscle & Nerve: Official Journal of the American Association of Electrodiagnostic Medicine. 1986;9(1):78–83.

13. Gossen ER, Ivanova TD, Garland SJ. Factors affecting the stability of the spike-triggered averaged force in the human first dorsal interosseus muscle. Journal of Neuroscience Methods. 2003 Jun 30;126(2):155–64.

14. Negro F, Yavuz UŞ, Farina D. Limitations of the Spike-Triggered Averaging for Estimating Motor Unit Twitch Force: A Theoretical Analysis. PLoS One. 2014 Mar 25;9(3):e92390.

15. Burke RE, Tsairis P. The Correlation of Physiological Properties with Histochemical Characteristics in Single Muscle Units. Annals of the New York Academy of Sciences. 1974;228(1):145–58.

16. Stephens JA, Usherwood TP. The mechanical properties of human motor units with special reference to their fatiguability and recruitment threshold. Brain research. 1977;125(1):91–7.

17. McNulty PA, Falland KJ, Macefield VG. Comparison of contractile properties of single motor units in human intrinsic and extrinsic finger muscles. The Journal of Physiology. 2000;526(2):445–56.

18. Lim KY, Rymer WZ, Thomas CK. Computational methods for improving estimates of motor unit twitch contraction properties. Muscle & Nerve: Official Journal of the American Association of Electrodiagnostic Medicine. 1995;18(2):165–74.

19. Raikova R, Pogrzebna M, Drzymała H, Celichowski J, Aladjov H. Variability of successive contractions subtracted from unfused tetanus of fast and slow motor units. Journal of Electromyography and Kinesiology. 2008;18(5):741–51.

20. Raikova R, Celichowski J, Pogrzebna M, Aladjov H, Krutki P. Modeling of summation of individual twitches into unfused tetanus for various types of rat motor units. Journal of Electromyography and Kinesiology. 2007;17(2):121–30.

21. Rohlén R, Raikova R, Stålberg E, Grönlund C. Estimation of contractile parameters of successive twitches in unfused tetanic contractions of single motor units – A proof-of-concept study using ultrafast ultrasound imaging in vivo. Journal of Electromyography and Kinesiology. 2022;67:102705.

22. Celichowski J, Raikova R, Aladjov H, Krutki P. Dynamic changes of twitchlike responses to successive stimuli studied by decomposition of motor unit tetanic contractions in rat medial gastrocnemius. Journal of Neurophysiology. 2014 Dec 15;112(12):3116–24.

23. Drzymała-Celichowska H, Celichowski J. Functional isolation of single motor units of rat medial gastrocnemius muscle. JoVE (Journal of Visualized Experiments). 2020;(166):e61614–e61614.

24. Calvin WH, Stevens CF. Synaptic Noise as a Source of Variability in the Interval between Action Potentials. Science. 1967;155(3764):842–4.

25. Negro F, Orizio C. Robust estimation of average twitch contraction forces of populations of motor units in humans. Journal of Electromyography and Kinesiology. 2017 Dec 1;37:132–40.

26. Negro F, Muceli S, Castronovo AM, Holobar A, Farina D. Multi-channel intramuscular and surface EMG decomposition by convolutive blind source separation. Journal of Neural Engineering. 2016;13(2):26027–26027.

27. Fuglevand AJ, Winter DA, Patla AE. Models of recruitment and rate coding organization in motor-unit pools. Journal of neurophysiology. 1993;70(6):2470–88.

28. Heckman CJ, Enoka RM. Physiology of the motor neuron and the motor unit. In: Handbook of Clinical Neurophysiology. Elsevier; 2004. p. 119–47.

29. Raikova RT, Aladjov HT. Hierarchical genetic algorithm versus static optimization—investigation of elbow flexion and extension movements. Journal of Biomechanics. 2002;35(8):1123–35.

30. Bonyadi MR, Michalewicz Z. Particle Swarm Optimization for Single Objective Continuous Space Problems: A Review. Evolutionary Computation. 2017 Mar 1;25(1):1–54.

31. Rohlén R, Lubel E, Farina D. Identification of motor unit discharges from ultrasound images: Analysis of in silico and in vivo experiments [Internet]. bioRxiv; 2024 [cited 2024 May 8]. p. 2024.01.18.576300. Available from: https://www.biorxiv.org/content/10.1101/2024.01.18.576300v1

32. Avrillon S, Hug F, Enoka R, Caillet AH, Farina D. The decoding of extensive samples of motor units in human muscles reveals the rate coding of entire motoneuron pools. eLife [Internet]. 2024 May 8 [cited 2024 Sep 27];13. Available from: https://elifesciences.org/reviewed-preprints/97085

33. Henneman E. Relation between size of neurons and their susceptibility to discharge. Science. 1957;126(3287):1345–7.

34. Henneman E, Somjen G, Carpenter DO. Excitability and inhibitibility of motoneurones of different sizes. Journal of Neurophysiology. 1965;28(3):599–620.

35. Holobar A, Farina D, Gazzoni M, Merletti R, Zazula D. Estimating motor unit discharge patterns from high-density surface electromyogram. Clinical Neurophysiology. 2009 Mar 1;120(3):551–62.

36. Del Vecchio A, Úbeda A, Sartori M, Azorín JM, Felici F, Farina D. Central nervous system modulates the neuromechanical delay in a broad range for the control of muscle force. Journal of Applied Physiology. 2018 Nov;125(5):1404–10.

37. Contreras-Hernandez I, Arvanitidis M, Falla D, Negro F, Martinez-Valdes E. Achilles tendon morpho-mechanical parameters are related to triceps surae motor unit firing properties. Journal of Neurophysiology [Internet]. 2024 Sep 4 [cited 2024 Oct 1]; Available from: https://journals.physiology.org/doi/abs/10.1152/jn.00391.2023

