## Supporting Information for "Model-based deconvolution of a force signal to estimate motor unit twitch parameters under low, moderate and high force isometric contractions"

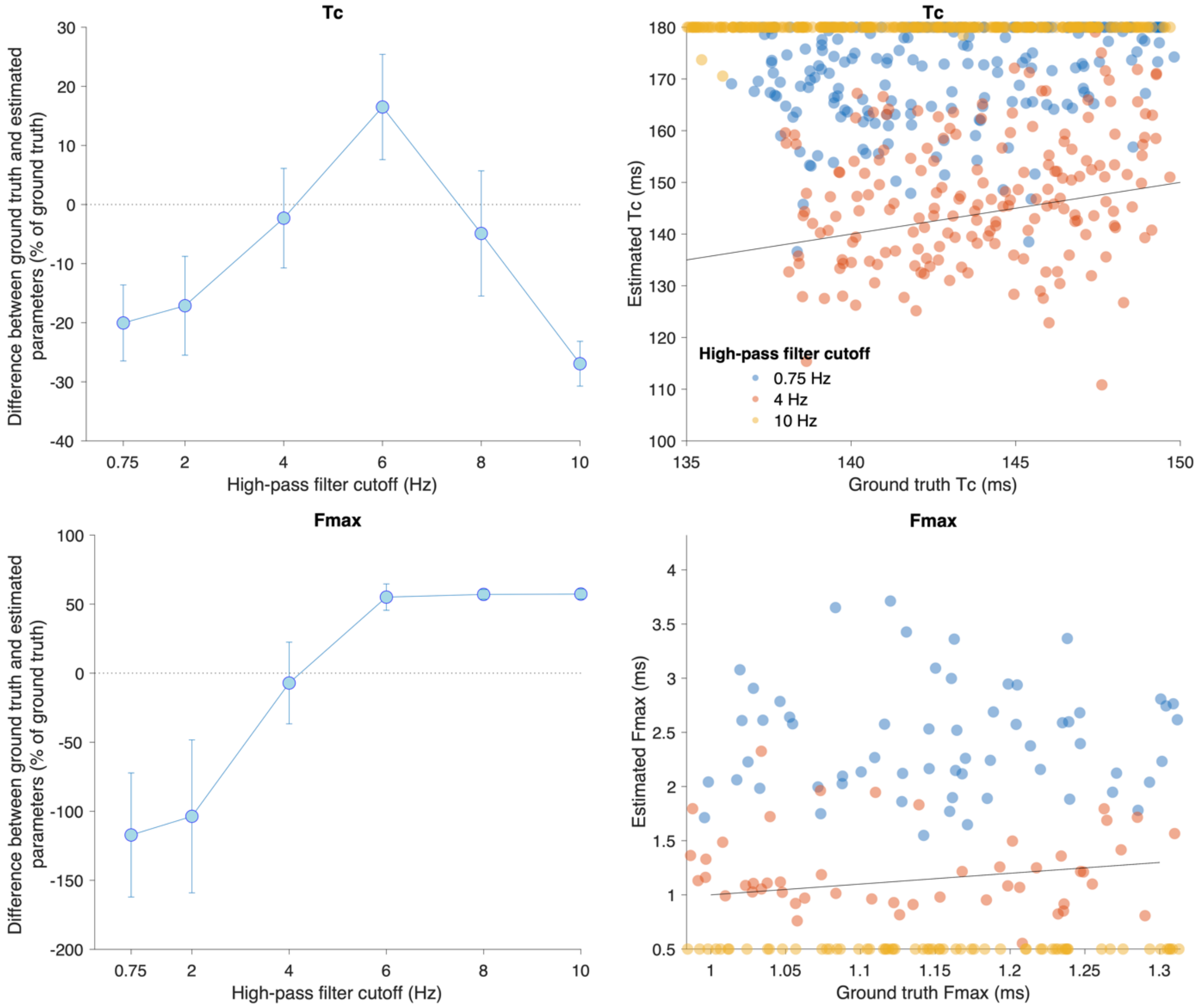


***Fig S1.*** *Finding the optimal high-pass filter cutoff by comparing the performance between ground truth and estimated parameters for simulations with a trapezoid excitatory drive function with 2.5% of the maximal excitatory drive. 4 Hz high-pass filter cutoff provides the less biased estimates.*

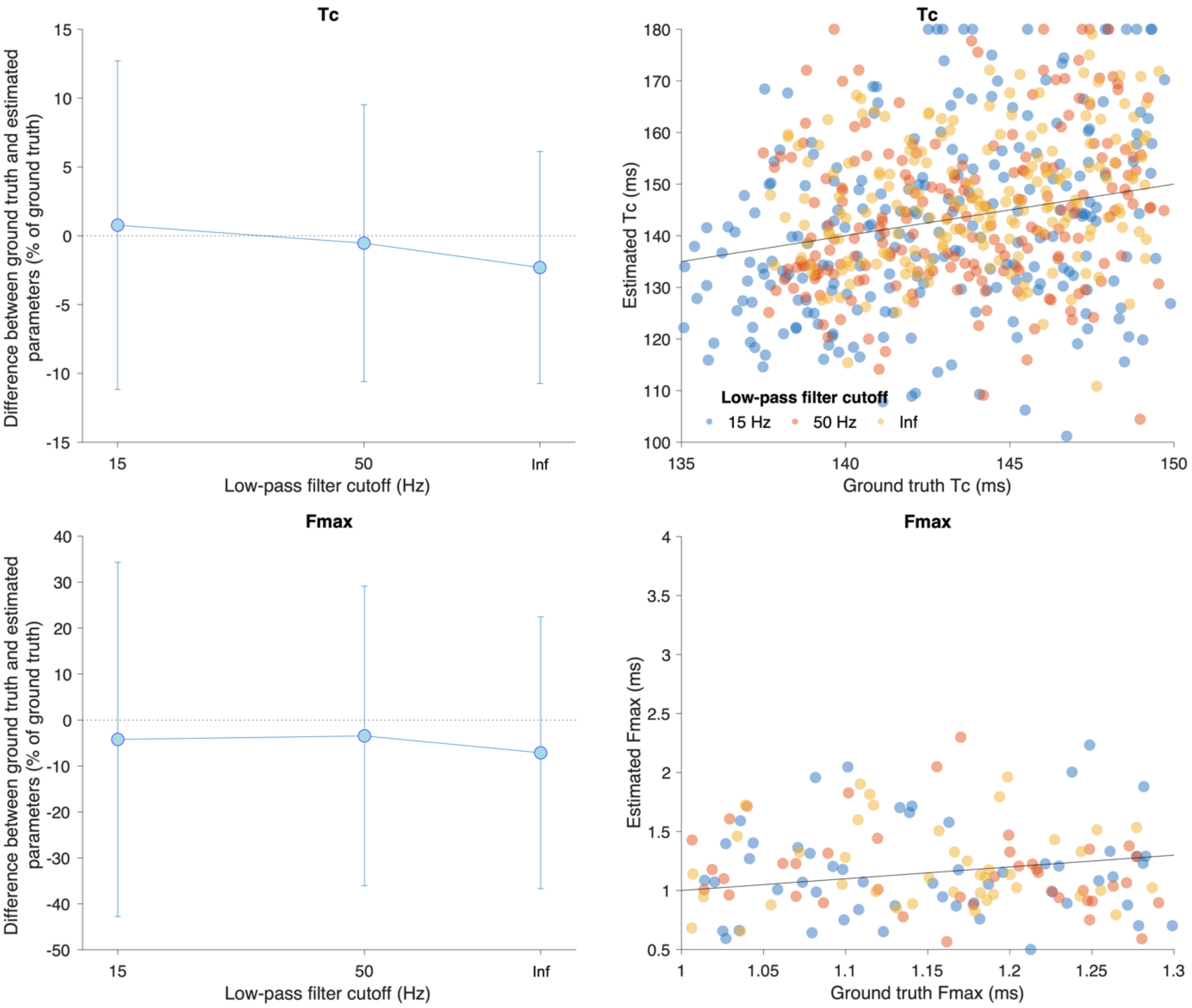


***Fig S2.*** *Finding the optimal low-pass filter cutoff by comparing the performance between ground truth and estimated parameters for simulations with a trapezoid excitatory drive function with 2.5% of the maximal excitatory drive. Using no low-pass filter provides the least variation.*

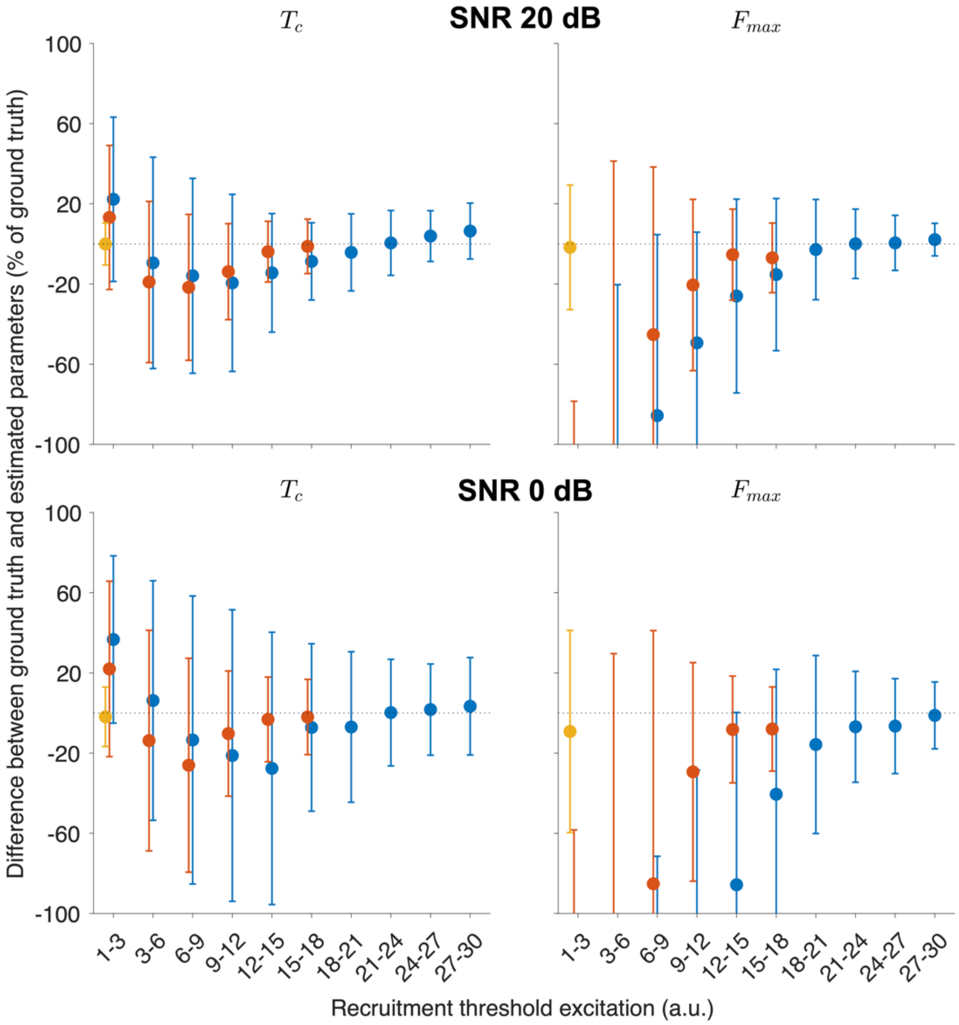


***Fig S3.*** *When the force signal has a lower signal-to-noise ratio (from 20 dB to 0 dB), the model-based deconvolution method provides estimates with a higher variation ($p < 0.001$), although still unbiased ($p > 0.05$). Note that this was without low pass filtering the force signal.*

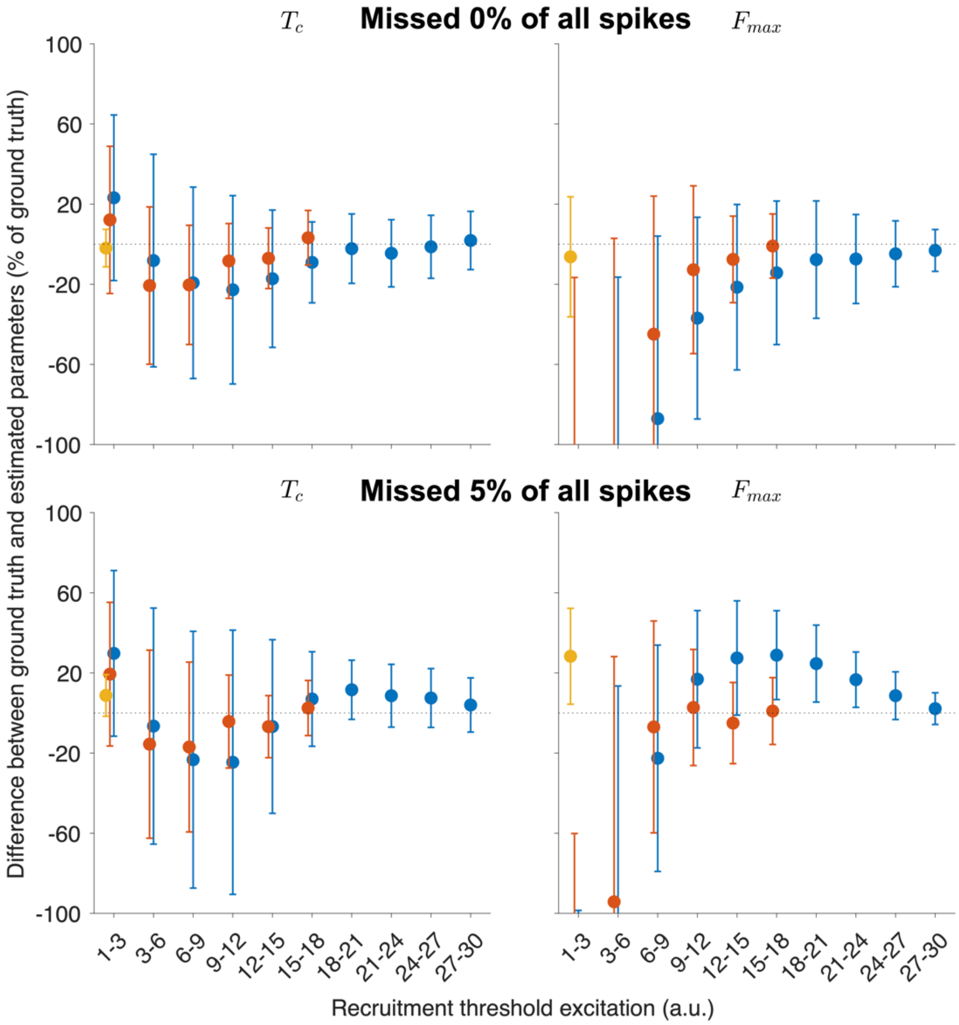


***Fig S4.*** *If the deconvolution method only uses 95% of the spikes (5% missed spikes), the estimate variation is the same, but there is an increased underestimation for the higher recruitment thresholds ($p < 0.001$).*

***Table S1.*** *Nonparametric Sign Test for the difference between the ground truth and estimated twitch parameters from the model-based deconvolution method for equal and unequal successive twitch profile cases. *** $p < 0.001$, ** $p < 0.01$, * $p < 0.05$.*

| | **Equal successive twitch case - Unbiasedness** | | | | | | | | | | | |
|---|---|---|---|---|---|---|---|---|---|---|---|---|
| | **2.5%** | | | | **30%** | | | | **70%** | | | |
| | **Tlead** | **Tc** | **Thr** | **Fmax** | **Tlead** | **Tc** | **Thr** | **Fmax** | **Tlead** | **Tc** | **Thr** | **Fmax** |
| **[1,3]** | 0.183 | * | * | 0.237 | * | *** | *** | *** | *** | *** | *** | *** |
| **[3,6]** | | | | | *** | *** | *** | *** | *** | 0.122 | *** | *** |
| **[6,9]** | | | | | *** | *** | *** | *** | 0.135 | *** | 0.189 | *** |
| **[9,12]** | | | | | * | *** | *** | *** | *** | *** | * | *** |
| **[12,15]** | | | | | 0.055 | *** | ** | ** | *** | *** | *** | *** |
| **[15,18]** | | | | | 1.000 | 0.690 | 0.690 | 1.000 | * | *** | *** | * |
| **[18,21]** | | | | | | | | | 0.154 | * | * | 0.393 |
| **[21,24]** | | | | | | | | | 0.600 | 0.600 | 0.162 | 0.485 |
| **[24,27]** | | | | | | | | | 0.439 | 1.000 | * | * |
| **[27,30]** | | | | | | | | | 0.098 | 0.410 | 0.784 | 0.410 |

| | **Unequal successive twitch case - Unbiasedness** | | | | | | | | | | | |
|---|---|---|---|---|---|---|---|---|---|---|---|---|
| | **2.5%** | | | | **30%** | | | | **70%** | | | |
| | **Tlead** | **Tc** | **Thr** | **Fmax** | **Tlead** | **Tc** | **Thr** | **Fmax** | **Tlead** | **Tc** | **Thr** | **Fmax** |
| **[1,3]** | *** | *** | *** | 0.294 | *** | *** | *** | *** | *** | *** | *** | *** |
| **[3,6]** | | | | | *** | *** | *** | *** | *** | * | *** | *** |
| **[6,9]** | | | | | *** | *** | *** | *** | * | *** | 0.787 | *** |
| **[9,12]** | | | | | *** | *** | *** | *** | *** | *** | *** | *** |
| **[12,15]** | | | | | *** | *** | *** | ** | *** | *** | *** | *** |
| **[15,18]** | | | | | * | *** | *** | 0.689 | *** | *** | *** | *** |
| **[18,21]** | | | | | | | | | *** | *** | *** | ** |
| **[21,24]** | | | | | | | | | * | ** | * | 0.750 |
| **[24,27]** | | | | | | | | | 0.456 | 0.351 | 0.709 | * |
| **[27,30]** | | | | | | | | | 1.000 | 1.000 | 1.000 | 1.000 |

***Table S2.*** *Nonparametric Sign Test for the difference between the ground truth and estimated twitch parameters from the spike-triggered averaging method for equal and unequal successive twitch profile cases.* *** $p < 0.001$, ** $p < 0.01$, * $p < 0.05$.

| | **Equal successive twitch case - Unbiasedness** | | | | | |
|---|---|---|---|---|---|---|
| | **2.5%** | | **30%** | | **70%** | |
| | **Tc** | **Fmax** | **Tc** | **Fmax** | **Tc** | **Fmax** |
| **[1,3]** | *** | *** | 0.350 | *** | *** | *** |
| **[3,6]** | | | *** | *** | *** | *** |
| **[6,9]** | | | *** | *** | *** | *** |
| **[9,12]** | | | *** | *** | *** | *** |
| **[12,15]** | | | *** | 0.665 | 0.578 | *** |
| **[15,18]** | | | *** | 0.424 | *** | *** |
| **[18,21]** | | | | | *** | *** |
| **[21,24]** | | | | | *** | *** |
| **[24,27]** | | | | | *** | *** |
| **[27,30]** | | | | | *** | ** |
| | **Unequal successive twitch case - Unbiasedness** | | | | | |
| | **2.5%** | | **30%** | | **70%** | |
| | **Tc** | **Fmax** | **Tc** | **Fmax** | **Tc** | **Fmax** |
| **[1,3]** | *** | *** | * | *** | *** | *** |
| **[3,6]** | | | 0.281 | 0.198 | *** | *** |
| **[6,9]** | | | *** | *** | *** | *** |
| **[9,12]** | | | *** | *** | *** | *** |
| **[12,15]** | | | *** | *** | 0.122 | *** |
| **[15,18]** | | | *** | *** | *** | *** |
| **[18,21]** | | | | | *** | *** |
| **[21,24]** | | | | | *** | *** |
| **[24,27]** | | | | | *** | *** |
| **[27,30]** | | | | | *** | * |
